# Ecogeographic signals of local adaptation in a wild relative help to identify variants associated with improved wheat performance under drought stress

**DOI:** 10.1101/2024.03.20.585976

**Authors:** Moses Nyine, Dwight Davidson, Elina Adhikari, Marshall Clinesmith, Huan Wang, Alina Akhunova, Allan Fritz, Eduard Akhunov

**Affiliations:** Department of Plant Pathology, Kansas State University, Manhattan, USA; Wheat Genetics Resource Center, Kansas State University, Manhattan, USA; Bayer, Chesterfield, USA; Syngenta, Junction City, USA; Broad Institute, Cambridge, Boston, USA; Integrated Genomics Facility, Kansas State University, Manhattan, USA; Department of Agronomy, Kansas State University, Manhattan, USA

## Abstract

Prioritizing wild relative diversity for improving crop adaptation to emerging drought-prone environments is challenging. Here, we combined the genome-wide environmental scans (GWES) in wheat diploid ancestor *Aegilops tauschii* with allele testing in the genetic backgrounds of adapted cultivars to identify new diversity for improving wheat adaptation to water-limiting conditions. Evaluation of adaptive allele effects was carried out in *Ae. tauschii*-wheat introgression lines (ILs) phenotyped for multiple agronomic traits under irrigated and water-limiting conditions using both UAS-based imaging and conventional approaches. The GWES showed that climatic gradients alone explain most (57.8%) of genomic variation in *Ae. tauschii*, with many alleles associated with climatic factors in *Ae. tauschii* being linked with improved performance of ILs under water-limiting conditions. The most significant GWES SNP located on chromosome 4D and associated with temperature annual range was linked with reduced canopy temperature in ILs. Our results suggest that (i) introgression of climate-adaptive alleles from *Ae. tauschii* have potential to improve wheat performance under water-limiting conditions, (ii) variants controlling physiological processes responsible for maintaining leaf temperature are likely among the targets of adaptive selection in a wild relative, and (iii) adaptive variation uncovered by GWES in wild relatives has potential to improve climate resilience of crop varieties.

## Introduction

The wild relatives of modern crops are a valuable source of adaptive diversity for developing improved varieties (Gill *et al*. 2006; Sohail *et al*. 2011; Kishii 2019). However, only a small fraction of wild relative diversity from the germplasm collections is utilized in breeding. The size of these collections, which may include thousands of accessions, complicates the selection of the most relevant genotypes for improving traits of interest. The prioritization of genebank germplasm for breeding climate adapted varieties is especially challenging due to the polygenic nature of adaptation to local environments (Araus *et al*. 2007; Exposito-Alonso *et al*. 2019). Therefore, the development of effective strategies, which are aimed at prioritizing wild relative germplasm for specific breeding applications, remains critical (Bohra *et al*. 2022).

The allopolyploid bread wheat, the second most important crop worldwide, originated by the hybridization of three wild grass species from the *Triticum* and *Aegilops* genera (Kihara 1944; Nesbitt and Samuel 1996; Dvorak *et al*. 1998; Tanno and Willcox 2006; Luo *et al*. 2007; Ozkan *et al*. 2011; Avni *et al*. 2017). Since its origin 10,000 years ago, wheat was disseminated by human migration and trade to diverse geographic regions with distinct climatic conditions(Balfourier *et al*. 2019). Archeological records and analyses of ancient DNA samples suggest that wheat reached Britain about 8,000 years ago (YA) (Smith et al., 2015) and China and Africa about 3,000 YA (Shewry 2009). Selection for performance in these diverse environments enriched local wheat populations for alleles contributing to adaptation to new climatic conditions (He *et al*. 2019; Zhao *et al*. 2023). However, the future climate change scenarios predict that climatic conditions in many wheat growing areas could be outside of the adaptive range of existing genotypes and lead to severe yield reduction (Tack et al., 2015; Ortiz-Bobea et al., 2019). The results of climate modeling suggest that nearly 40% of crop growing areas might require new varieties to sustain crop production (Schlenker and Roberts 2009; Zabel et al., 2021). Leveraging the genetic diversity of multiple wild ancestors of wheat, which are evolved to grow in diverse environments, is one of the promising strategies for broadening the climate adaptive potential of modern wheat.

The direct ancestors of wheat, *Aegilops tauschii* and *Triticum turgidum* ssp. *dicoccoides* (wild emmer), are two most broadly distributed species among the wheat wild relatives (Dvorak *et al*. 1998; Avni *et al*. 2017). These species are also among the most represented in germplasm collections, some of which host thousands of accessions of *Ae. tauschii* and *T. turgidum* (Sharma *et al*. 2021). Because these wild ancestors of wheat share homologous genomes, their chromosomes could easily recombine, facilitating introgression of allelic diversity from *Ae. tauschii* and wild emmer into wheat (Nyine *et al*. 2020). By using the synthetic hexaploid wheat (SHW) lines, which are hybrids of tetraploid wheat and *Ae. tauschii*, the allelic diversity of these ancestors was introduced into multiple international breeding programs from CIMMYT, ICARDA, China, Australia, United Kingdom, and United States (Pestsova et al., 2004; Börner et al., 2015). For example, it was demonstrated that these ancestors of wheat have potential to improve adaptation to water limiting conditions and heat, and increase biomass and harvest index (Singh et al., 2019; Molero et al., 2023). However, considering the broad adaptive potential of these species reflected in their wide geographic distribution, the question remains of how effective these efforts were at capturing adaptive diversity of *Ae. tauschii* and wild emmer.

The prioritization of wild relative accessions for pre-breeding of climate resilient crops remains challenging. Wild relatives could be phenotypically pre-screened for target traits. However, this screening could be performed only for simple traits, and has limited utility for complex adaptive traits if phenotypic evaluation was not performed in the genetic background of adapted cultivars. Another approach to prioritize accessions is the development of “core collections” assembled from a large number of genotypes selected to maximize the genetic diversity of the sample (Frankel 1984). While this strategy could effectively reduce the number of accessions, its major disadvantage in application to large collections is that it targets only common adaptive alleles, removing rare alleles or allelic complexes. The third approach is based on selection of wild relative accessions based on environmental parameters at the site of accession’s origin (Turner et al., 2010; Jones et al., 2012; Lasky et al., 2015) (Bari *et al*. 2012). However, while this strategy could capture alleles contributing to an adaptive phenotype of small number of accessions, its ability to maximize the recovery of adaptive genetic diversity at species-wide level would be limited.

A combination of genome diversity analyses with the geographic patterns of environmental variation for detecting adaptive diversity is another strategy that so far had limited usage in crop breeding. The cost-efficiency of next-generation sequencing (NGS) genotyping approaches made possible generating genome-wide variation for geographically diverse populations. By combining genomic data with eco-geographic variables, it became possible to identify alleles associated with adaptive phenotypes. These approaches, referred to as genome-wide environmental scans (GWES), identify loci involved in local adaptation based on a high correlation between allele frequencies and eco-geographic variables. In an early GWES study, a number of climate-associated alleles (CAA) were mapped in *Arabidopsis* by using 13 climatic variables, among others including extremes and seasonality of temperature and precipitation (Hancock et al., 2011). The CAA identified by GWES allowed for accurate prediction of the relative fitness of *Arabidopsis* accessions in local environments (Turner et al., 2010; Hancock et al., 2011; Frachon et al., 2018). The GWES in sorghum and Mexican white oak detected adaptive variants that also produced reliable phenotypic predictions (Lasky et al., 2015; Martins et al., 2018). These studies suggest that adaptive alleles identified using the GWES have potential to predict agronomic phenotypes in target environments.

Though GWES were shown to be effective at identifying loci contributing to environmental adaptation, it remains unclear whether these loci could be used to prioritize wild relative accessions for introgression into modern crop varieties to improve their adaptive potential in extreme environments. To address this question, we used a diverse collection of *Ae. tauschii* to conduct GWES and identified variants contributing to climatic adaptation. We specifically focused on those variants that correlate with precipitation and temperature gradients during growth season. Then, we selected a geographically diverse set of *Ae. tauschii* accessions to develop introgression populations by crossing them with the adapted wheat varieties. The developed introgression population was grown for several seasons across diverse environments and agronomic performance of introgression lines (ILs) was assessed by measuring agronomic and physiological traits. The physiological status and growth of ILs were evaluated using the UAS-based phenotyping with the RGB and thermal cameras. The wheat productivity was assessed by measuring yield and yield component traits (thousand grain weight, grain area, grain width and grain length). The relationship between phenotypic data and climate adaptive alleles introgressed from *Ae. tauschii* was investigated to better understand the value of GWES in wild relatives as a tool for selecting wild relative accessions to improve the adaptive potential of wheat varieties.

## Materials and methods

### Plant materials

A diverse set of 137 geo-referenced *Ae. tauschii* accessions collected over a geographic range of species distribution and representing locations with diverse historic climatic and bioclimatic characteristics was acquired from the USDA NSGC to identify the CAAs (Table S1). A subset of 21 geographically diverse accessions was selected from this population and crossed with hard red winter wheat varieties to generate *Ae. tauschii-*wheat amphiploids. The amphiploids were then crossed with six hard red winter wheat cultivars adapted to grow in the US Great Plains to develop *Ae. tauschii-*wheat ILs (Nyine et al., 2020, Nyine et al., 2021). A total of 351 BC_1_F_3:5_ introgression lines that had phenology similar to that of the recurrent parents were used to study the impact of introgressed CAA on the adaptative traits.

### Genotyping and imputation

DNA was extracted from two-week old seedling leaf tissues of the diverse *Ae. tauschii* accessions and the derived introgression population using DNeasy 96 Plant DNA extraction kit (Qiagen) following the manufacturer’s protocol. The quality and concentration of the DNA was assessed using PicoGreen dsDNA assay kit (Life Technologies). The extracted DNA was normalized to 400 ng (20ul of 20ng/ul) using the Qiagility robot (Qiagen). Genotyping by sequencing (GBS) included a library size selection step performed using the Pippin Prep system (Sage Scientific) to enrich the library for 270-330 bp fragments, as described in Saintenac et al. (2013). The prepared libraries were sequenced on Illumina NextSeq 500. Variant calling was done using the TASSEL v5.0 GBS v2 pipeline (Glaubitz et al., 2014).

To increase the density of SNP markers in both populations, we re-sequenced the panel of 21 *Ae. tauschii* accessions and the six recurrent hexaploid wheat lines using whole-genome sequencing approach. PCR-free genomic libraries were constructed using Illumina protocol at the Integrated Genomic Facility (IGF) at Kansas State University. Paired-end sequences (2 x 150 bp) were generated using NovaSeq at Kansas University Medical Center and NextSeq 500 at IGF. The data were combined and processed as described by Nyine et al. (2021). Missing and ungenotyped SNPs in the *Ae. tauschii* diversity panel and the introgression population were imputed from the parental genotypes using Beagle v5.0 (Browning and Browning 2013). After imputation and filtering out SNPs with genotype probability below 0.7, we retained 6,365,631 SNPs in *Ae. tauschii* diversity panel and 5,208,054 SNPs in the introgression population.

### Population structure and variance partitioning of SNP diversity in *Ae. tauschii*

To understand the level of genetic diversity within the *Ae. tauschii* population and how both geography and climate shaped the SNP variation in the population, we pruned the 6.3 million SNPs based on linkage disequilibrium (LD) using PLINK v1.9 and retained 109,627 SNPs that had r^2^ < 0.5 in 50 kb sliding window with step size of 5 kb. The proportion of ancestry shared between accessions was estimated from the LD pruned SNPs and the geographical coordinates for the accessions’ collection sites using the tess3r R package (Caye et al. 2016). The maximum number of ancestral populations tested was eight (K = 1:8). Each K was run 10 times for 200 iterations (rep = 10, max.iteration = 200) and the spatial projection of ancestral coefficients was based on least squares method (method = “projected.ls”). The optimal number of ancestral populations selected based on cross-validation scores was K = 4 because it split the *Ae. tauschii* population into two lineages and four sub-lineages that coincided with previous findings by Wang et al., (2013). A plot showing the population admixture and the spatial distribution of accessions from different subspecies at the sites of sample collection was generated.

The proportion of SNP variance in the *Ae. tauschii* accessions was partitioned into those explained by geographic distance and climate using the ‘varpart’ function in R package ‘vegan’. The geographic distances were calculated using the ‘distVicentyEllipsoid’ function in R package ‘geosphere’ using the GPS coordinates from the accessions’ collection sites and the results were presented in a Venn diagram.

### Redundancy analysis

The diverse set of *Ae. tauschii* accessions used in this study came from a wide range of geographical locations with distinct climatic and bioclimatic conditions suggesting that certain genetic factors are involved in local adaptation. Based on this hypothesis we modeled the relationship between response variables (SNPs) and explanatory variables (climatic and bioclimatic factors, and geographic distance) using the redundancy analysis (RDA) (Van den Wollenberg 1977; Lasky et al., 2015) and mixed linear models to identify variants contributing to local adaptation. Variables were ranked to identify those that contribute most to SNP diversity in the *Ae. tauschii* population. To achieve this, the ordiR2step function was applied on the RDA results with adjusted R^2^ using the forward selection method from 10,000 permutations. Type 1 error was minimized during the selection of the most important factors contributing to SNP diversity by following the rules proposed by Blanchet et al., (2008). The full RDA model based on all climatic and bioclimatic variables with the calculated adjusted R^2^ values was evaluated to determine variables that (1) significantly improved the explained variation of SNP diversity distribution in *Ae. tauschii* population at alpha 0.05, and (2) whose total adjusted R^2^ did not exceed the adjusted R^2^ value of the full model. Based on the aforementioned conditions, the most important variables identified were projected on the first two principal components as a biplot. To illustrate the variation of temperature annual range (BIO7) from the *Ae. tauschii* accessions’ collection sites, a heatmap was plotted using the ‘heat_point’ function provided in R package ‘autoimage’ and an overview of the variation of the most important variables at the collection sites for the sub-lineages was compared using boxplots. All variables were scaled to range between 0 and 1 by dividing with the highest value within the dataset and then squaring them to eliminate the negatives before generating the boxplots with ggplot2 in R.

### Identification of climate associated alleles (CAAs) in *Ae. tauschii*

To determine the genetic basis of local adaptation in *Ae. tauschii*, we used both RDA and GWAS to identify SNPs that were significantly associated with geographic, climatic and bioclimatic variables. We extracted the first three RDA loadings for each SNP from the RDA model described above and transformed them to Z-scores. The mean Z-score was calculated from all SNPs and any SNP with three standard deviations from the mean was considered a candidate CAA SNP. Pearson’s correlation coefficients were used to determine the variable with the strongest association to each SNP, thus each CAA SNP was assigned to only one variable. To capture most CAA, we also performed a GWAS using a compressed mixed linear model in GAPIT (Lipka et al., 2012). The geographic, climatic and bioclimatic variables were used as phenotypes. The population structure was accounted for by including PCAs calculated from the marker data as covariates in the model. Multiple test correction was performed using the Benjamini-Hochberg’s method (FDR ≤ 0.05).

### Phenotyping of *Ae. tauschii* introgression population

The population of ILs was phenotyped under field conditions for three seasons between 2018 and 2020 to evaluate the adaptive potential of CAAs introgressed in the winter wheat. In 2018 and 2019, phenotyping was done at Colby (Kansas, USA) under irrigated and non-irrigated conditions. In 2020, phenotyping was done at Ashland (Kansas, USA) under non-irrigated conditions. The experimental layout at all locations followed an augmented design with six recurrent hexaploid wheat parents and three additional winter wheat lines adapted to Kansas weather as controls. Experimental plots were 2.5 m x 0.5 m consisting of three rows separated by 18 cm. During planting, granular 18-46-0 diammonium phosphate (DAP) fertilizer was applied at a rate of 168.1 kg/ha and liquid 28-0-0 urea ammonium nitrate (UAN) was applied at a rate of 67.3 kg/ha in the spring to supply additional nitrogen to the plants. The lateral irrigation system was used to maintain the soil moisture in the irrigated block.

The ILs were phenotyped for yield and the component traits such as spikelet number per spike (SNS), thousand grain weight (TGW), grain area (GA), grain width (GW) and grain length (GL). During the growing season, remote sensing data including RGB, NDVI and canopy temperature (CT) were collected at multiple time points during growth seasons using unmanned aerial system (UAS) mounted with specific sensors for each data type to evaluate the physiological status and growth trend of the introgression lines. The RGB and NDVI imagery data were processed in Agisoft software (version) to generate orthomosaics and digital elevation models (DEM) whereas the thermal data were processed in Pix4D to generate the CT orthomosaics. The raster files generated by Agisoft and Pix4D were imported into QGIS v3.4 software for plot level data extraction. Shape files consisting of rectangular polygons that overlaid each plot in the experimental block were created and the mean pixel values for each color band within the polygon were calculated using raster zonal statistics tools and saved as a comma separated values (csv) file. Other indices such as visible atmospherically resistant index (VARI) and triangular greenness index (TGI) were derived from the RGB data whereas NDVI was derived from near infrared and red color bands using the following equations:

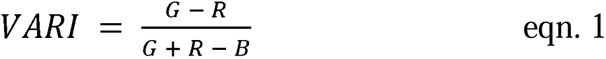

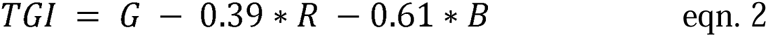

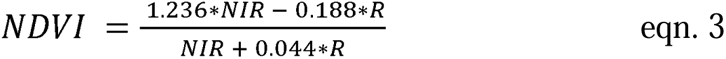

where R, G, B and NIR are the mean pixel values for the red, green, blue and near infrared color bands.

Heading data were collected in 2020 at Ashland and validated in 2022 at RockyFord, Manhattan, Kansas USA. Heading date was recorded when 50 % of spikes fully emerged from the flag leaf. The number of days to heading (DTH) were calculated by subtracting the planting date from the heading date. To understand how much the heading date for the introgression lines varies from the controls, we calculated the mean DTH for the controls and subtracted the DTH for each introgression to generate the deviation in days to heading (DDTH).

### Best linear unbiased predictions (BLUPs)

Best linear unbiased predictions for yield and yield component traits were obtained from a mixed linear model implemented in R package. Given that the experimental layout followed an augmented design all controls were given a code 1, and the test introgression lines were assigned a 0. Each introgession line was assigned a unique numeric code which was used as a group identifier in all experiments whereas the controls were assigned a 999 regardless of the accession as the group identifier. A mixed linear model was run for different traits. For example, BLUPs for the number of days to heading, used in GWAS analysis were estimated from the following model:

DTH∼Loc+Check, random= ∼ Acc + Acc:range/row + Acc:LocName,

where DTH is number of days to heading, Loc is the field trial location, Check defined lines whether they are controls or test lines and Acc is the accessions.

Canopy temperature data were collected over multiple time points (aka flights) in the two years. Spatial correction was performed using SpATS implemented in MrBean, a shiny based R package. Variance due to genotype and environment were estimated as well as narrow sense heritability. After excluding outliers, BLUEs were predicted for the test lines based on the variance in the controls. All flights and blocks with heritability less than 0.2 were excluded from the linear mixed model. The remaining data were used to estimate genotype BLUPs across flights, treatment blocks and years. In the model, experimental treatment and flights were considered as fixed effects whereas genotypes were considered as random effects.

### Association between CAAs and phenotypic traits in introgression population

The frequency spectra of CAAs derived from *Ae. tauschii* was estimated for groups of ILs that have trait values falling into the tails of phenotype distributions. Our expectation was that if the CAAs are associated with variation in phenotypic traits in the introgression population, the phenotypic value tails should show CAA frequency spectra distinct from the CAA frequency spectra for the whole population. For this purpose, we ranked the ILs based on trait values and compared the CAA frequency spectrum in the whole population (WP) and the lower and upper 5th percentile of the phenotype distributions. Lines were considered to belong to the lower or upper tail groups if they were ranked as outliers in at least two trials. The frequency spectrum was built using 5,675 CAAs in the introgression population by counting alleles in 9 allele frequency bins.

Allele frequency of all CAA sites was calculated for the whole population and for the introgression lines that ranked in the lower and upper 5% tails of phenotype distribution using vcftools. Differentiation in allele frequency was determined by calculating the fold change (FC) in allele frequency between the introgression lines in the tails and the whole population. SNPs with FC ≥ 2 were considered to strongly differentiated. For some traits, where the allele frequency FC was less than two across all sites, the threshold was adjusted accordingly.

To determine the relationship between recombination rate gradient and allelic differentiation, we split the chromosome arms into three equal parts. The number of differentiated CAAs within each chromosome segment was counted using the ‘bedmap’ function of BEDOPs tools. The redundant CAA sites showing the high levels of LD were removed using PLINK. We retained only those SNPs that had r^2^ < 0.5 within the 50 kb window with a step size of 5 kb. The total number of differentiated alleles was aggregated for all traits and chromosomes.

To confirm the contribution of CAAs to adaptation traits a linear regression of yield on CT was performed to determine the proportion of variance in yield explained by the variation in CT. Introgression lines that ranked in the 5th and 95th percentiles of CT distribution were compared for yield performance relative to the recurrent parents. GWAS was performed on the traits phenotyped in the BC_1_F_3:5_ *A. tauschii*-wheat introgression population including CT, heading date, yield and component traits to determine loci with significant associations. Multiple GWAS models were tested on each trait with varying number of principal components to correct for population structure. GWAS analysis was implemented in GAPIT v3.0. CAAs significantly associated with traits in the introgression population at the FDR value 0.05 were considered adaptive in the winter wheat background.

## Results

### Environmental scans in *Aegilops tauschii*

The genetic basis of *Ae. tauschii* adaptation to diverse climatic conditions across a broad geographic range extending from Eastern Europe to China remains poorly understood. To identify genetic loci contributing to adaptation, we conducted genotype-environment association analyses using 109,627 SNPs identified in a geographically diverse panel of 137 accessions. This set of SNPs was selected by LD-based pruning from a larger set including 6,365,631 genotyped and imputed SNPs. Using this data, we explored the population structure of our samples and its correspondence to the previously identified four main lineages (L1W, L1E, L2E, L2W) of *Ae. tauschii* (Wang et al. 2013) (Fig. 1A). The inferred population structure of *Ae. tauschii* accessions was consistent with the results of previous studies (Wang et al. 2013), showing the split between L1 and L2 lineages, where L1 was composed of *Ae. tauschii* ssp. *tauschii* and L2 included accessions of *Ae. tauschii* ssp. *strangulata*, the closest ancestor of the wheat D genome (Wang et al. 2013), (Fig. S1A). The first two principal components, separating L1 and L2 lineages, accounted for 78.4% of the variation in our samples (Fig. S1B). The split between the L1W and L1E and between the L2E and L2WE was also obvious in our panel.

**Fig. 1.**
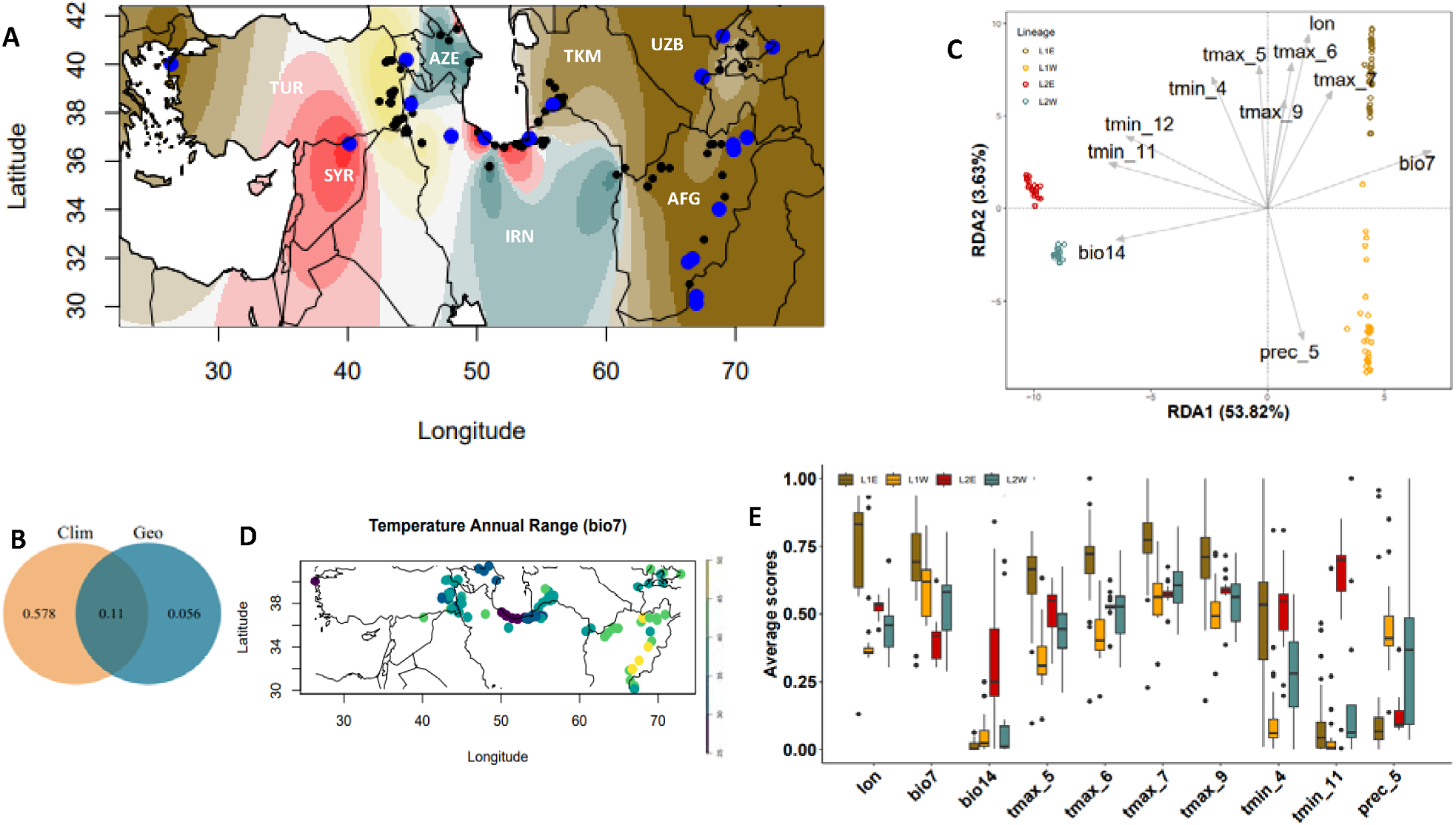
Ecogeographic distribution of 137 diverse *Ae. tauschii* accessions. **A**) Map shows the geographic locations and the ancestry coefficients of 137 accessions. Twenty-one accessions used to develop the *Ae. tauschii*-wheat introgression population are shown in blue. **B)** Proportion of SNP variance explained by climate (Clim) and geographical distance (Geo) between accessions. **C)** Redundancy analysis (RDA) plot showing the 11 best explanatory variables for SNP variance in *Ae. tauschii* accessions including temperature annual range (bio7), precipitation of driest month (bio14), precipitation in May (prec_5), minimum temperature in April, November and December (tmin_4, tmin_11 and tmin_12, respectively), maximum temperature in May, June, July and September (tmax_5, tmax_6, tmax_7 and tmax_9, respectively) and longitude (lon). **D)** Heatpoint map showing the variation in temperature annual range (bio7) at the sampling locations of the *Ae. tauschii* accessions. **E)** Boxplots showing the variation in lon, bio7, bio14, tmax_5, tmax_6, tmax_7, tmax_9, tmin_4, tmin_11 and prec_5 in the ecogeographic locations of different *Ae. tauschii* lineages. Lineage 1 East (L1E) and Lineage 1 West (L1W) belong to *Ae. tauschii* ssp. *tauschii* whereas Lineage 2 East (L2E) and Lineage 2 West (L2W) belong to *Ae. tauschii* ssp. *strangulata*.

To identify variants contributing to local adaptation, we modeled the relationship between response variables (SNPs) and explanatory variables (climatic and bioclimatic factors, and geographic distance) using redundancy analysis (RDA) (Van den Wollenberg 1977; Mcardle and Anderson 2001; Lasky et al. 2015). For this purpose, we used historical data for the bioclimatic and climatic factors estimated for the geographic locations at the accession collection sites. The total SNP variance explained by both geographic distance and climatic and bioclimatic factors was 85.2% (Fig.1B), with the adjusted R^2^ value being 69.6%. Climate alone accounted for 57.8% of the SNP variation in *Ae. tauschii*, whereas geographic distance between accessions and the interaction between geographic distance and climate accounted for 5.6% and 11% of SNP variation, respectively. These results indicate that the distribution of SNP variation among *Ae. tauschii* accessions is primarily driven by gradient in climatic and bioclimatic factors rather than by *Ae. tauschii* geographic dispersal.

Depending on their impact on adaptive traits, individual climatic factors could have distinct effects of SNP variation among accessions (Hancock et al., 2011; Lasky et al., 2015; Li et al., 2021; Chang et al., 2022). The first two RDAs accounted for 22.58% of the total SNP variation in the population. A triplot with two RDAs shows separation of the population into four distinct groups (Fig. S1C) coinciding with the previously detected split between the L1E, L1W, L2E and L2W subpopulations. Among the geographic variables, longitude and altitude showed the strongest effect on SNP distribution between the two subspecies of *Ae. tauschii* followed by latitude (Fig. S1D). The environmental variables contributing most to SNP variation were determined using the ‘ordiR2step’ function in R package ‘vegan’ using the forward selection method and 10,000 permutations (Blanchet et al., 2008). A total of 11 variables were detected, including temperature annual range (bio7), precipitation of driest month (bio14), precipitation in May (prec_5), minimum temperature in April, November and December (tmin_4, tmin_11 and tmin_12, respectively), and maximum temperature in May, June, July and September (tmax_5, tmax_6, tmax_7 and tmax_9, respectively) and longitude (lon) (Table 1, Fig. 1C). The tmin_11, tmin_12, bio7, and bio_14 contributed most to RDA1 that explains most of the genetic differentiation between the two subspecies of *Ae. tauschii*. The prec_5, tmin_4, tmax_5, tmax_6, tmax_7 and tmax_9 factors contributed to RDA2 that explains most of the genetic differentiation between the Eastern (L1E, L2E) and Western (L1W, L2W) populations of the two *Ae. tauschii* lineages. These results indicate that temperature and precipitation gradients during the growth periods coinciding with flowering, grain filling and maturation were the main factors that shaped SNP diversity in *Ae. tauschii* and likely contributed to genetic differentiation among the four lineages.

**Table 1.** The geographic, climatic and bioclimatic variables that contributed most to SNP variation in *Ae. tauschii*. The R^2^.adj are cumulative values.

| Variable | $R^2$.adj* | Df | AIC | F | Pr(>F) | |
| --- | --- | --- | --- | --- | --- | --- |
| Temperature annual range (bio7) | 0.27664 | 1 | 1236.8 | 43.4509 | 1.00E-04 | *** |
| Min temperature in November (tmin_11) | 0.35655 | 1 | 1224.7 | 14.6606 | 1.00E-04 | *** |
| Max temperature in July (tmax_7) | 0.39818 | 1 | 1218.2 | 8.5394 | 0.0008 | *** |
| Max temperature in June (tmax_6) | 0.43887 | 1 | 1211.3 | 8.832 | 0.0011 | ** |
| Min temperature in December (tmin_12) | 0.49921 | 1 | 1199.5 | 13.8934 | 0.0002 | *** |
| Longitude (lon) | 0.52234 | 1 | 1195.1 | 6.1326 | 0.0043 | ** |
| Max temperature in September (tmax_9) | 0.53963 | 1 | 1191.9 | 4.9424 | 0.0093 | ** |
| Precipitation of driest month (bio14) | 0.55291 | 1 | 1189.6 | 4.0891 | 0.0148 | * |
| Min temperature in April (tmin_4) | 0.57148 | 1 | 1185.7 | 5.4646 | 0.004 | ** |
| Max temperature in May (tmax_5) | 0.59765 | 1 | 1179.6 | 7.6331 | 0.0017 | ** |
| Precipitation in May (prec_5) | 0.60705 | 1 | 1177.8 | 3.4162 | 0.025 | * |

Two subspecies of *Ae. tauschii*, ssp. *strangulata* and ssp. *tauschii* appear to show different levels of adaptation to distinct climatic conditions. We compared the distribution of the main climatic and geographic factors (lon, bio7, bio14, tmax_5, tmax_6, tmax_7, tmax_9, tmin_4, tmin_11, tmin_12 and prec_5) between the two subspecies of *Ae. tauschii* (Fig.1E). Analysis of variance showed significant differences between the accessions from these subspecies (Table 2). Results suggest that L1E lineage is adapted to warmer and drier conditions of Eastern Iran, Afghanistan, Turkmenistan, Uzbekistan, Tajikistan and Kyrgyzstan (Table S1), indicating that these *Ae. tauschii* accessions could be a good source of drought and heat stress tolerance. The lowest precipitation of driest month characterized by high maximum temperature from May up to September was one of the major differentiating ecogeographic factors for L1E. In contrast, L1W is represented by accessions mostly from Eastern Turkey and Northwestern Iran where a high precipitation is recorded in May and significantly lower maximum temperature from May to September. The factors that contribute most to the genetic differentiation of this sublineage are tmin_4, tmax_5 and prec_5.

**Table 2.** Comparison of geographic and climatic factors between the main sub-lineages of *Ae. tauschii*.

| Variable | Lineage 1 East<br>(L1E) | Lineage 1<br>West (L1W) | Lineage 2 East<br>(L2E) | Lineage 2 West<br>(L2W) |
| --- | --- | --- | --- | --- |
| lon | 62.83<sup>a</sup> | 45.98<sup>c</sup> | 52.69<sup>b</sup> | 49.09<sup>bc</sup> |
| bio7 | 40.08<sup>a</sup> | 37.01<sup>b</sup> | 30.11<sup>c</sup> | 34.88<sup>b</sup> |
| bio14 | 3.13<sup>c</sup> | 7.10<sup>b</sup> | 19.90<sup>a</sup> | 10.16<sup>b</sup> |
| tmax_5 | 269.44<sup>a</sup> | 195.90<sup>c</sup> | 244.45<sup>b</sup> | 221.77<sup>b</sup> |
| tmax_6 | 324.80<sup>a</sup> | 251.92<sup>b</sup> | 283.60<sup>b</sup> | 273.94<sup>bc</sup> |
| tmax_7 | 345.08<sup>a</sup> | 295.97<sup>b</sup> | 300.45<sup>b</sup> | 305.41<sup>b</sup> |
| tmax_9 | 292.64<sup>a</sup> | 253.33<sup>c</sup> | 272.45<sup>b</sup> | 262.71<sup>bc</sup> |
| tmin_4 | 80.82<sup>a</sup> | 32.77<sup>c</sup> | 84.80<sup>a</sup> | 53.00<sup>b</sup> |
| tmin_11 | 18.26<sup>b</sup> | 0.03<sup>c</sup> | 78.25<sup>a</sup> | 28.65<sup>b</sup> |
| tmin_12 | -17.46<sup>b</sup> | -49.28<sup>c</sup> | 42.35<sup>a</sup> | -23.00<sup>b</sup> |
| prec_5 | 26.72<sup>c</sup> | 58.36<sup>a</sup> | 30.60<sup>c</sup> | 46.41<sup>b</sup> |

Means with the same superscript letters are not significantly different. The following factors were considered: longitude (lon), temperature annual range (bio7), precipitation of driest month (bio14), maximum temperature in May, June, July and September (tmax\_5, tmax\_6, tmax\_7 and tmax\_9), minimum temperature in April, November and December (tmin\_4, tmin\_11 and tmin\_12) and precipitation in May (prec\_5). Statistical significance is based on Tukey's honestly significant difference test at 95% confidence level.

Generally, *Ae. tauschii* ssp. *strangulata* lineages are found around the Caspian Sea with some accessions found in Syria and Turkey. L2E is adapted to a relatively uniform precipitation and mild temperature which are characteristic of the Southern Caspian Sea in Northern Iran. The L2W accessions are mostly found near Western Caspian Sea in Azerbaijan and parts of Northwestern Iran. The region is characterized by moderate to high variation in climatic and bioclimatic factors. Amongst the main variables differentiating L2W from other sublineages are tmin_4 and prec_5 (Table 2).

### Mapping adaptive SNPs in *Ae. tauschii*

The results of both redundancy analysis (RDA) and genome-wide association mapping (GWAS) were used combined to identify SNPs that are significantly associated with variation in geographic, climatic and bioclimatic variables across the *Ae. tauschii* sampling locations. The first three RDA loadings for each SNP were extracted from the RDA model, transformed to Z-scores, and climate associated alleles (CAA) were defined as outlier SNPs with three standard deviations from the mean Z-score. After removing duplicate SNPs, a total of 10,149 D-genome SNPs showed a positive correlation (mean = 0.58, range 0.14-0.82) with 51 out of 58 variables analyzed in our study (Table S2, Table S3).

Temperature annual range (bio7) was ranked as the most significant variable accounting for 27.66% of the SNP variation in *Ae. tauschii* population based on the adjusted R^2^ values (P < 0.001). It had the highest number of correlated SNPs (747) amongst other most significant variables. Chromosomes 1D and 7D had the highest number of SNPs showing significant correlation with the geographic, climatic and bioclimatic variables (Fig. 2A). Genome-wide association mapping is another approach that was previously used for studying genome-by-environment interactions (Wallace et al. 2016). By using a compressed mixed linear model, we identified a total of 10,569 D-genome SNPs significantly associated with 42 out of 58 variables (Table S2, Fig. 2B). Most of these variants were located on chromosomes 1D and 5D (Fig. 2A and Table S4). Unlike in RDA analysis, where each SNP was assigned to a single highly correlated variable, in GWAS, many SNPs were associated with more than one variable at FDR ≤ 0.05. Combined, RDA and GWAS identified 18,096 SNPs with significant association to geographic, climatic and bioclimatic variables (Table S5). Among the SNPs identified, a set of 2,622 SNPs were detected using both methods (Fig. 2C, Table S6). The functional annotation of these SNPs using SnpEff (Cingolani et al., 2012) showed that only 29 of them were stop codon gain, missense, synonymous, intronic or splice region variants (Table S6). The majority of SNPs (2,316) were intergenic variants, and 277 SNPs were located 5 kb upstream or downstream of gene models.

**Fig. 2.**
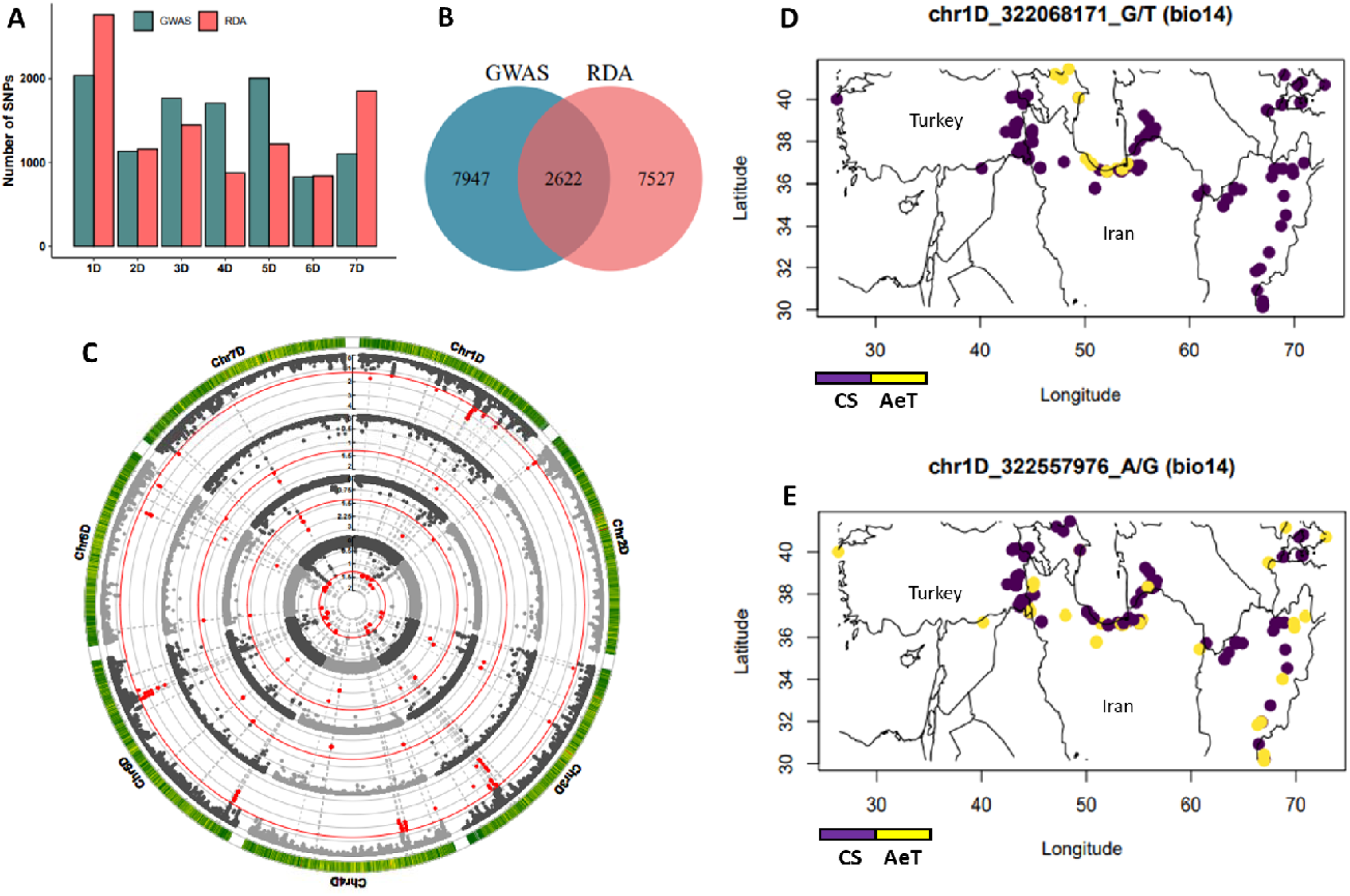
Number of climate associated SNPs per chromosome identified by the redundancy analysis (RDA) and genome-wide association analysis (GWAS), and the geographical distribution of SNP alleles associated with precipitation of driest month (bio14). A) Chromosome distribution of SNPs identified through RDA and GWAS that were associated with different geographic, climatic and bioclimatic variables. B) Venn diagram showing the total number of climate associated SNPs identified by RDA and GWAS. C) Circular Manhattan plot showing GWAS for four of the most significant variables. Starting from the innermost circle outward are minimum temperature in April, maximum temperature in May and June (tmax_5 and tmax_6) and bio14. The red lines show an FDR threshold of 0.05 and the red dots are the significant SNPs on each chromosome. D) *Ae. tauschii* (AeT) specific allele (yellow) on chromosome 1D showing adaptation to areas with high precipitation of driest month near the Caspian Sea. E) *Ae. tauschii* specific allele (yellow) on chromosome 1D showing adaptation to areas with a wide range of reduced precipitation in the driest month. The purple color shows the reference allele similar to Chinese Spring (CS).

Consistent with prior studies, the geographic extent of CAAs could primarily be explained by the distribution of climatic factors (Hancock et al., 2011). For example, among SNPs significantly associated with bio14, there is an *Ae. tauschii* allele (chr1D_322068171) that was found only in accessions from the region near the Caspian Sea (Fig. 2D), which shows a high precipitation of driest month (Table 1). Another bio14-associated SNP (chr1D_322557976) had an allele identified in accessions from a broad geographic region characterized by low precipitation of driest month experiencing extreme drought stress due to high temperature from May up to September (Fig. 2E, Table 1). These results suggest that *Ae. tauschii* could be the source of adaptive alleles to a broad range of climatic factors useful for addressing the impact of climate change on wheat productivity.

### Evaluation of the adaptive potential of *Ae. tauschii* CAAs in winter wheat

To evaluate the ability of CAAs from *Ae. tauschii* to improve the adaptive potential of bread wheat, we developed a wild relative introgression population using a set of 21 diverse accessions that were selected to capture the ecogeographical and allelic diversity of species (Nyine et al., 2020; Nyine et al., 2021). To facilitate comparison with parental lines, ILs in the populations were selected to match development and phenology of hexaploid wheat parents (Nyine et al., 2020). Out of 18,096 climate adaptive SNPs identified by RDA and GWAS, 31.4% (5,675 CAA SNPs) were present in the introgression population. It is likely that loss of some of the CAAs in introgression population could be caused by their linkage with deleterious alleles selected against during population development (Nyine et al., 2020) (Fig. 3). Among the introgressed CAAs, a total of 1,089 SNPs were detected using both environmental association scan methods.

**Fig. 3.**
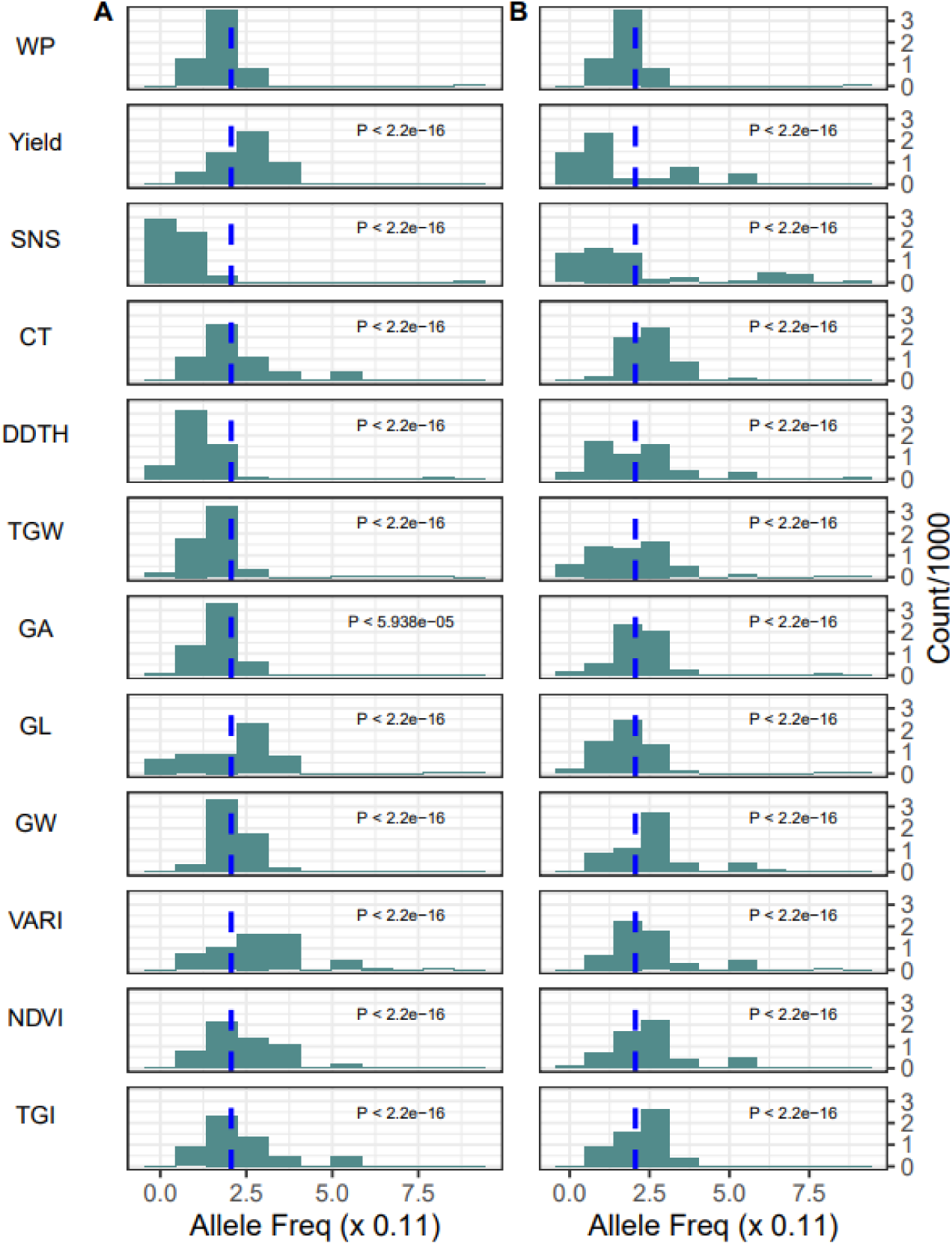
Frequency spectra of climate associated alleles (CAA) in the tails of phenotype distribution relative to the allele frequency spectra in the whole population (WP). The A and B panels show the 5^th^ and 95^th^ percentiles of the yield, spikelet number per spike (SNS), canopy temperature (CT) deviation in days to heading (DDTH) from the control mean DTH, thousand grain weight (TGW), grain area (GA), grain length (GL), grain width (GW), visible atmospherically resistant index (VARI), normalized difference vegetation index (NDVI) and triangular greenness index (TGI) traits.

Introgression of beneficial alleles occurs in both high and low recombining regions of the genome. While introgressions found in high recombining regions become shorter after a few generations of recombination, those in the low recombining regions tend to persist as large linkage blocks. The large introgression blocks in the pericentromeric regions of the chromosome could have unintended consequences on non-targeted traits due to linkage with deleterious alleles linked to adaptive SNPs and epistatic interactions with adapted genetic background. When selection is applied, the frequency of adaptive alleles in the high recombining regions usually increases whereas the frequency of adaptive alleles in low recombining regions may (i) increase if SNP effect on an adaptive trait is stronger than the combined negative effects of linked alleles or (ii) reduce if the combined negative effects of linked alleles are stronger than the effects of adaptive SNPs. In a breeding population, shifts in allele frequency are best observed in the tails of distributions for phenotypes targeted by selection. We compared the frequency spectra of CAAs derived from *Ae. tauschii* in the tails of phenotype distributions in the introgression population. Our expectation was that if CAAs affect wheat performance, the tails of distribution for yield and yield component, CT and DDTH traits should show distinct frequency spectra. For this purpose, we ranked the ILs based on trait values and compared the CAA frequency spectrum in the whole population (WP) with CCA frequency spectra in the lower and upper 5th percentile tails of the phenotype distribution. The frequency spectra were generated for the 5,675 CAAs in the introgression population by counting *Ae. tauschii* alleles in the 9 allele frequency bins. Lines in the tails of the trait distribution were filtered to retain only those that ranked in the same percentile group for at least two traits.

A significant shift from the mean CAA frequency (0.226) in the WP was observed in the tails of phenotype distribution for various traits (Fig 3). The shift in the CAA frequency spectrum in the tails of yield distribution was significantly different from WP mean (Kolmogorov-Smirnov test: lower tail P < 2.2e-16; upper tail P < 2.2e-16). In the lower tail of yield distribution, the frequency of CAAs was high suggesting that lines with large introgression segments that had many CAAs that are likely in LD with deleterious alleles contributed to yield penalty. The top yielding lines showed a bimodal distribution of CAA frequency, suggesting the occurrence of both negative and positive selection at different CAA loci in these lines. For SNS however, a decrease in the *Ae. tauschii* allele frequency was linked with low spikelet number per spike whereas a combination of both low and high frequency *Ae. tauschii* alleles were associated with with higher SNS (Kolmogorov-Smirnov test: lower tail P < 2.2e-16; upper tail P < 2.2e-16).

Previous studies have shown a positive relationship between yield and SNS, especially if all spikelets are fertile and produce seeds (Rawson 1970; Zhang et al., 2018, Kuzay et al., 2019). However, yield is a complex trait modulated by changes heading date and yield component traits, such as grain area, width and length. The shift in the CAA frequency spectrum for the deviation in days to heading (DDTH) distribution from the WP mean followed the same pattern as that observed for SNS (Kolmogorov-Smirnov test: lower tail P < 2.2e-16; upper tail P < 2.2e-16) suggestive of the shared biological pathways between these two traits. These results are as expected because longer development period and hence delayed heading have been associated with increase in SNS (Rawson 1970). Guo et al. (2018) attributed the increase in spikelet number to delayed spikelet initiation and transition from double-ridge phase to terminal spikelet which coincided with delayed heading date.

Canopy temperature (CT) is one of the critical physiological traits that reflects the adaptive potential of plants in local environments (Kumar et al., 2017) (Still *et al*. 2021) and could be used to identify drought and heat stress tolerant plant genotypes. Previous studies demonstrated that CT in wheat is a complex quantitative trait mostly linked to QTLs that control root architecture necessary for improved water use efficiency and maintenance of transpiration rate (Pinto and Reynolds 2015). While in the low CT tail, most CAAs had lower than average allele frequency, we detected some CAAs that significantly increased in frequency compared to population mean suggestive of their contribution to regulation of CT. CT negatively correlated with yield under drought stress (r = –0.45, P = 0.0) which agreed with the previous studies that showed the importance of CT depression for increasing yield in wheat (Pinter et al. 1990; Amani et al. 1996). Introgression from *Ae. tauschii* into spring wheat was associated with low CT and improved yield under heat stress (Molero et al., 2023). Previously, we showed that the difference in mean yield of some *Ae. tauchii* introgression lines in our population reached 57% when compared to the checks under drought stress conditions (Nyine et al., 2021).

Besides CT, both visible atmospherically resistant index (VARI) and normalized difference vegetation index (NDVI) are correlated vegetation indices used to monitor plant health and biomass accumulation. Lines in both lower and upper tails showed a significant shift towards high frequency CAAs. The latter shows that some CAAs could be associated with the positive impact on vegetation indices and physiological status of the plants under stress. The finding of CAAs showing strong shift in the lower tails of both traits suggest that some CAAs or linked introgression variants could be associated with the negative impact on these traits. The triangular greenness index (TGI) is an indicator of total chlorophyll content in the leaves (Hunt et al., 2013) which is useful for estimating the stay green characteristics in wheat (Lopes and Reynolds 2012). In this population, lines that matured late were those with the highest number of *Ae. tauschii* alleles thus the TGI values were also high during the growing season.

Both natural and artificial selection results in genetic differentiation at target loci between the selected and non-selected populations. Since the efficiency of selection at target loci is higher in the high recombining regions, for CAAs linked with trait variation, we expect to observe higher frequency differentiation between the lines in phenotypic tails in the high-recombining terminal regions of chromosomes rather than in the low-recombining pericentromeric regions. By plotting CAAs identified for eleven most significant climatic and bioclimatic variables along the chromosomes, we show that differentiated CAAs are enriched in the high-recombing regions of chromosomes (Figs. 4C and 4D). These results suggest that 1) selection of lines in the phenotypic extremes of agronomic and physiological traits prioritizes those that carry CAAs located within the high-recombining regions of the genome likely due to the reduced linkage to deleterious alleles, and 2) introgressed CAAs are associated with variation in phenotypic traits linked with wheat performance in both irrigated and water-limiting conditions.

**Fig. 4.**
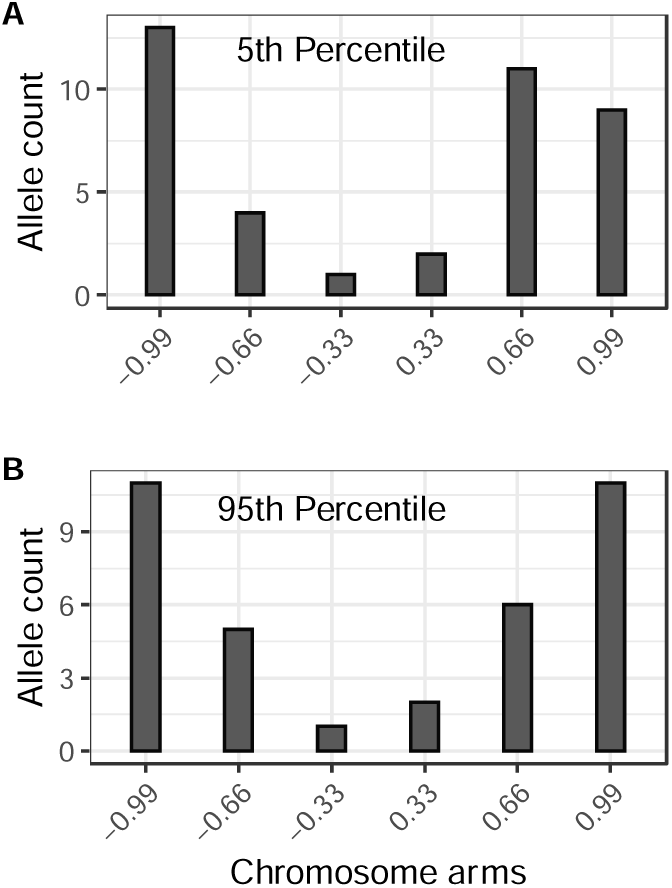
Chromosome distribution of CAAs differentiated between the tails of phenotypic extremes in introgression population. The CAAs detected for most significant climatic and bioclimatic variables were included into the analyses. Each chromosome arm was split into three regions with each region representing 33.3% of arm length. The counts of CAAs in each region across all chromosomes in the wheat genome was combined. The A and B panels show chromosome distributions for differentiated CAAs in the 5^th^ and 95^th^ percentile tails of phenotype distribution, respectively.

### CAAs are linked with variation in adaptive traits in the introgression population

Association analyses were performed in the introgression population using a set of 5,675 climate-adaptive SNPs to determine variants that contribute to improved performance of introgression lines under the water-limiting and irrigated conditions. The CT, heading date, yield and yield component traits were used as plant performance metrics. Significant CAA-trait associations with heading date, spikelet number per spike, grain width and length were observed in the introgression population (Supplementary File S1). Considering the importance of CT for assessing the physiological response of plants to drought stress (Pinto and Reynolds 2015; Kumar et al., 2017), we focused on the results of association analyses between CAAs and CT.

The CT data were collected using unmanned aerial system (UAS)-based thermal imaging from both irrigated and non-irrigated field trials at multiple time points during the growing season in 2018 and 2019. Variation in CT was influenced by both genotype and environment (Table 3). In the 2018 growing season, narrow sense heritability (*h***^2^**) for CT varied between 0.54 and 0.85 whereas in 2019 it ranged from 0.24 to 0.78. In the 2019 growing season, residual variance in CT was much higher than due to genotype effect. This could be linked to the fact that in 2019, Colby experienced high precipitation and low temperature conditions during the growing season. Best linear unbiased predictors for spatially corrected CT varied from –0.57 to 1.8 suggesting that some introgression lines were able to lower CT compared to others that had higher CT (Figure 5A). A comparison of yield and CT showed a strong negative relationship with CT accounting for 30% of yield variation in the *Ae. tauschii* introgression population (Fig. 5B). This result was confirmed by performing phenomic predictions using the random forest model with CT and yield component traits as predictors of yield (Fig. S2). These analyses showed that CT is the most significant factor for predicting yield followed by grain length and thousand grain weight in this population, consistent with previous observations (Wardlaw et al. 1989).

**Fig. 5.**
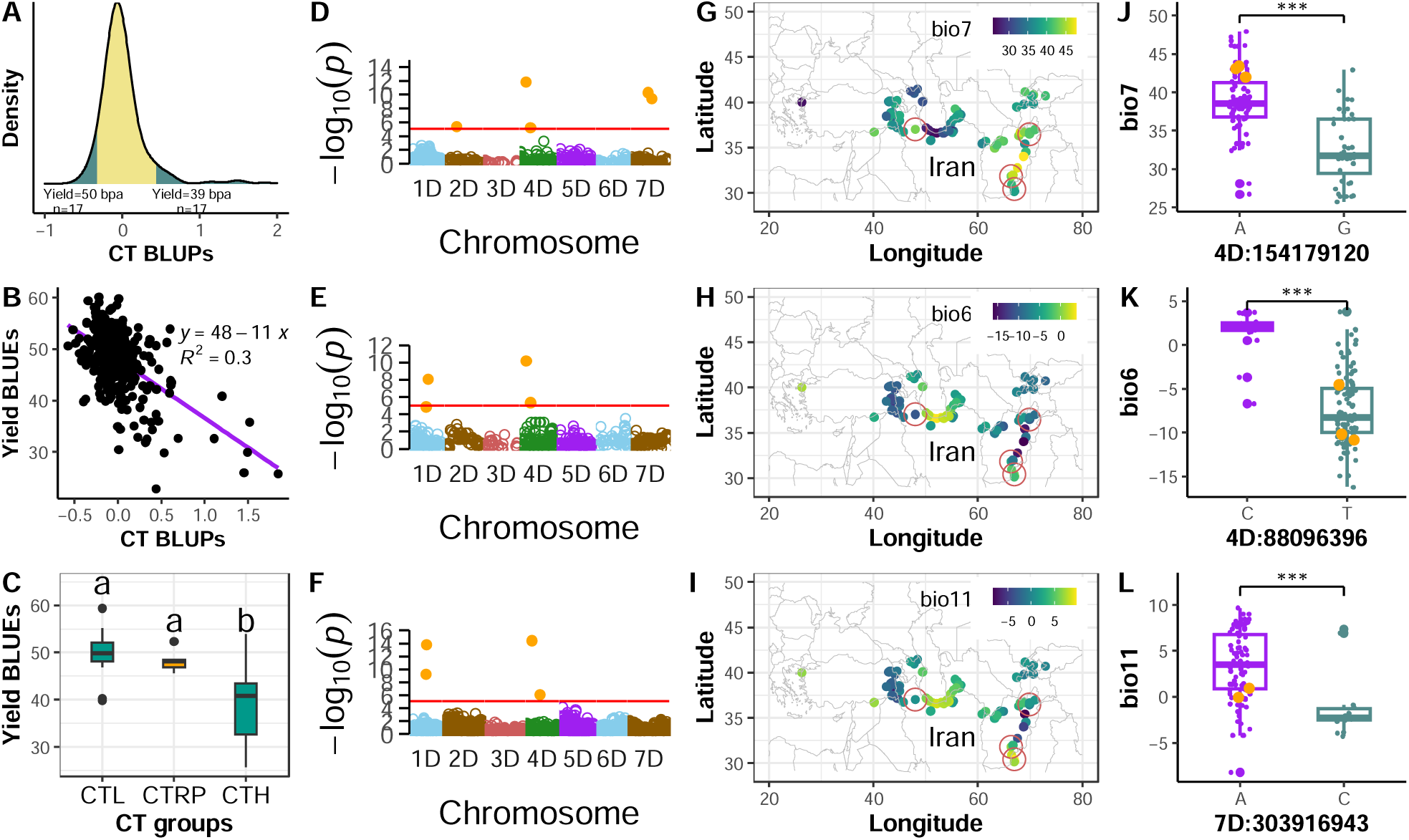
Relationship between **c**anopy temperature (CT) and yield performance of the introgression lines, the genomic loci associated with CT and origin of *Ae. tauschii* providing CT lowering alleles in hexaploid wheat background. (A) CT distribution for the introgression population at Colby in 2018 and 2019 growing seasons. The shaded tails represent the 5th and 95th percentiles. (B) Regression of yield on CT, (C) Yield of ILs showing low CT (CTL) and high CT (CTH) relative to recurrent parents CT (CTRP). Different letters on top of the box plots indicate significant differences at 95% confidence level, (D-F) Manhattan plots showing the quantitative trait nucleotides (QTN) associated with CT. Where D and E are based CAA SNPs with MLMM and BLINK models, respectively and F is based LD pruned SNPs with MLMM. The red line in the Manhattan plot indicates the P-value corresponding to a threshold FDR 0.05. (F-H) Heatpoint maps showing temperature annual range (bio7), min temperature in the coldest month (bio6) and mean temperature in coldest quarter (bio11) in the geographic origin of *Ae. tauschii* accessions. Accessions in red circles are the parents for introgression lines with low CT. (I-K) Distribution of bioclimatic variables (bio7, bio6 and bio11) in geographical origin of 137 *Ae. tauschii* accessions. Orange dots represent *Ae. tauschii* accessions used to generate introgression lines that rank in the 5th percentile for CT distribution.

**Table 3:**
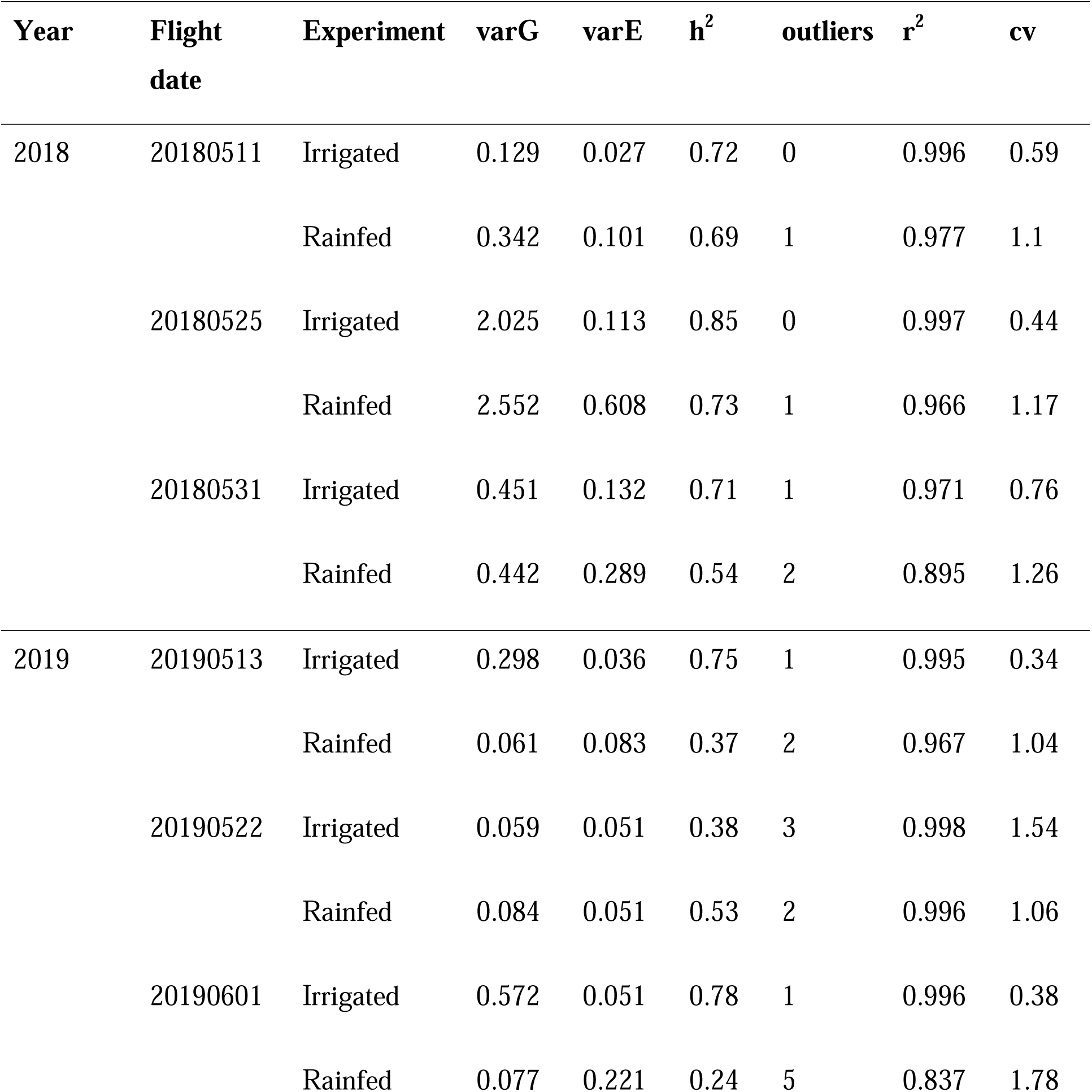
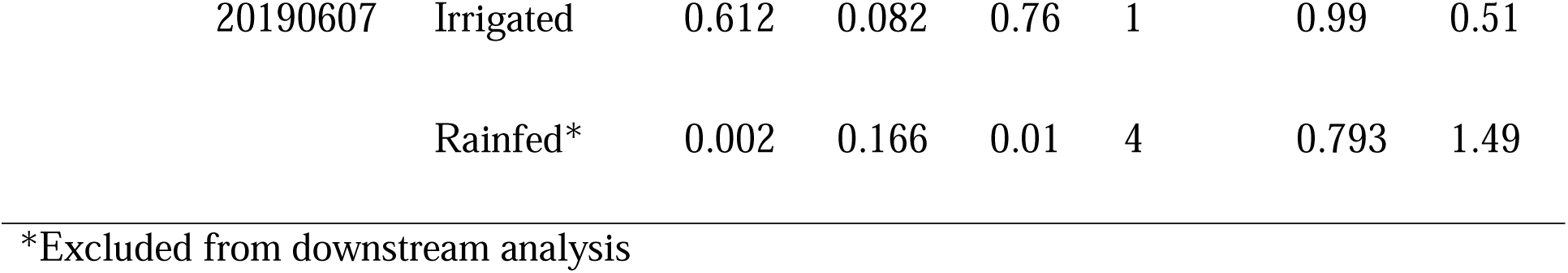
Effect of genotype and environment on canopy temperature variation and its heritability in the *Ae. tauschii* introgression population.

To further understand the impact of introgressed alleles from *Ae. tauschii* into hexaploid wheat background on yield, we identified ILs in the 5^th^ and 95^th^ percentiles of CT distribution and compared them to recurrent parents. The average yield for ILs in the 5^th^ percentile of CT was 50 bpa, which was higher but not significantly different from the recurrent parents (48 bpa, P = 0.825). The lack of significant difference could be attributed to high variation in yield in ILs showing low CT, suggesting that other genetic factors could contribute to final yield. The introgression lines in the 95^th^ percentile however, showed significant yield reduction (39 bpa) relative to the recurrent parents (Tukey HSD, P < 0.008), confirming the importance of CT trait for predicting grain yield in wheat.

To identify the genomic loci associated with CT depression, we performed GWAS using genotypes at CAAs (Segura et al., 2012; Wang and Zhang 2021). The multiple-locus mixed linear model (MLMM) and Bayesian-information and Linkage-disequilibrium Iteratively Nested Keyway (BLINK) revealed significant SNP-trait associations on chromosomes 1D, 2D, 4D and 7D (Fig. 5D, Table S7). Some of the most significant SNPs associated with CT on chromosomes 1D and 4D (chr1D_265243957 and chr4D_154179120) showed the highest correlation with temperature annual range (bio7, r = 0.59). Based on the RDA analyses, bio7 was identified as the most significant variable shaping SNP variation in *Ae. tauschii*. Other SNPs significantly associated with CT in introgression population correlated with the minimum temperature in the coldest month (bio6, r = 0.67), mean temperature in the coldest quarter (bio11, r = 0.6), longitude and precipitation in the driest months (August to October) (Table S7).

Besides using only SNP sites with CAAs for GWAS, 5.3 million SNPs from *Ae. tauschii* introgression population were pruned based on LD resulting in 99,529 SNPs with r^2^ ≤ 0.5. When GWAS was performed using this set of SNPs, the MLMM model revealed three QTLs that were associated with CT, including one on chromosome 1D and two on 4D. The significant SNPs on 1D were chr1D_254646871 and chr1D_265399733 (Fig 5F, Table S7). Although these SNPs were not part of the SNP set detected in the environmental scans of *Ae. tauschii* accessions, they were located within the same genomic intervals identified by genome-wide association mapping in the introgression population using the climate-associated SNPs. Within the interval 254 – 266 Mb, there were 191 CAA SNPs (Table S3) that highly correlated with mean diurnal range (bio2), temperature annual range (bio7), mean temperature of driest quarter (bio9), minimum temperature in March (tmin_3), precipitation in September (prec_9) and longitude (lon). Similarly, the first QTL on 4D contains SNP chr4D_92689640 (Table S7), and in the interval 90 – 94 Mb on 4D, four CAA SNPs identified in *Ae. tauschii* accessions were strongly correlated with bio7. The second QTL contains SNP chr4D_229986737 (Table S7), and a search for CAA SNPs 5 Mb to the left and right of the significant QTN identified 31 SNPs that were correlated with prec_9, tmin3 and lon variables.

The source of alleles lowering CT in the introgression lines were mostly from *Ae. tauschii* ssp. *tauschii* accessions (TA2388, TA2536, TA2521 and TA10177), collected from areas such as Afghanistan, Iran and Pakistan known for high bio7 (43), on average with nearly no precipitation in the driest quarter of the year (Figs. 5G, 5J). These results suggest that, *Ae. tauschii* growing in high temperature and low precipitation conditions could improve wheat adaptation to water-limiting conditions when introgressed into adapted wheat background.

## Discussion

The lineages of *Ae. tauschii* are spread over a large geographic area with a wide range of variation in climatic factors, including some of the locations with extremely dry and hot environments (Dvorak *et al*. 1998; Wang *et al*. 2013; Gaurav *et al*. 2022). The existence of strong SNP-climate correlations reported here provides effective means for detecting climate adaptive variants in diploid *Ae. tauschii* using environmental genome scans. The range of geographic distribution for adaptive variants varied broadly with some alleles showing narrow geographic distribution, and other alleles showing broad distribution across large geographic areas. Consistent with prior studies, these spatial patterns of allele distribution can primarily be explained by the distribution of climatic factors (Hancock et al., 2011) (Lasky *et al*. 2012, 2015). In our analyses the environmental and bio-climatic factors alone accounted for a substantial proportion (57.8%) of spatial genetic variation in *Ae. tauschii* with relatively small contribution from geographic dispersal (5.6%). These results suggest consistent environmental gradients across the *Ae. tauschii* distribution range likely shaped the spatial structure of genomic variation in this wild ancestor of wheat and contributed to genetic differentiation between its main lineages.

Correlations between genomic diversity and climatic factors indicate that the temperature and precipitation gradients during the growth periods coinciding with flowering, grain filling and maturation contributed to genetic differentiation among the four main lineages of *Ae. tauschii* and its two subspecies, *strangulata* and *tauschii*. The lowest precipitation in driest month was one of the major differentiating ecogeographic factors for L1E, whereas temperature and precipitation gradients in April and May contributed most to the genetic differentiation of L1W lineage. The variation in precipitation and temperature in April and May were among the main ecogeographic factors explaining differentiation between the L2W and L2E lineages of *Ae. tauschii* ssp. *strangulata*. The lineage L2E of *Ae. tauschii* ssp. *strangulata,* which contributed 10,000 years ago to the origin of bread wheat (Wang et al., 2013; Luo et al., 2017), grows in a narrow geographic region south of the Caspian Sea with limited variation in climatic factors characteric of the humid mild subtropical environments. As a result, adaptive diversity captured by the D genome of bread wheat is primarily restricted to those alleles that are represented in this region. The limited levels of gene flow detected between wheat and *Ae. tauschii* ssp. *strangulata* did not have dramatic impact on the genetic diversity of the D genome (Wang *et al*. 2013; He *et al*. 2019; Zhou *et al*. 2020; Gaurav *et al*. 2022). Thus, the polyploidization bottleneck associated with wheat origin resulted in not only the overall loss of genetic diversity in the wheat D genome (He et al., 2019; Gaurav et al., 2022) but also in the massive loss of adaptive alleles represented in all four sublineages of *Ae. tauschii*. While the consequences of the loss of these alleles in wheat are hard to predict, we might expect that it had a negative impact on the adaptive potential of hexaploid wheat and offset progress with development of drought-resilient wheat varieties.

Introgression from *Ae. tauschii* into hexaploid wheat had positive effects on traits playing an important role in increasing crop productivity and improving adaptation to drought. In our previous study, we showed that 3.2% of introgression lines carrying *Ae. tauschii* haplotypes outperformed parental lines in drought trials (Nyine et al., 2021). Consistent with these results, several high-yielding drought tolerant cultivars have been derived from synthetic wheat lines created using *Ae. tauschii* as one of the parents (Rosyara et al. 2019; Molero et al., 2023; Pinto and Reynolds 2015). Our analyses suggest that improved performance of wheat introgression lines could be largely attributed to introduction of climate-adaptive alleles that show association with environmental variation in *Ae. tauschii*. Statistically significant shifts in allele frequency in the extreme tails of phenotypic trait distributions and significant associations detected for climate-adaptive alleles in GWAS for canopy temperature and productivity traits support this conclusion.

The signatures of adaptation detected by environmental scans in the genome could be driven by complex historic gradients of environments and associated with diverse adaptive mechanisms (Lasky *et al*. 2012; Anderson and Song 2020). As a result, it is difficult to establish relationships between adaptive alleles from environmental scans and specific phenotypic traits measured for introgression populations in field trials. The temperature annual range (bio7) was among the main bio-climatic factors that contributed most to shaping the spatial genomic variation in *Ae. tauschii*. The SNP locus located on chromosome 4D showing strongest association with bio7 in environmental scans also showed strongest association with variation in canopy temperature in introgression lines. This result indicates that among the targets of selection imposed by variation in bio7 are variants associated with pathways controlling physiological processes responsible for maintaining canopy temperature under drought stress (Jackson *et al*. 1981). In the field trails, lines carrying *Ae. tauschii* alleles at loci associated with reduced canopy temperature were among the top yielding introgression lines suggesting that these *Ae. tauschii* alleles improve adaptation to drought stress and likely act as the main drivers of increased yield. Likewise, detection of *Ae.tauschii* introgression into chromosome 6D associated with reduction in canopy temperature and increased yield under dry conditions, confirms the importance of this adaptive mechanism for drought tolerance (Molero et al., 2023) (Still *et al*. 2021). These results indicate that environmental scans focusing on the relevant bio-climatic variables are an effective means for uncovering variants in wild relatives to improve wheat adaptation to water-limiting conditions and increase its yield potential.

Our study shows that whole genome sequencing of diverse collections of wild relatives integrated with environmental scans could provide an effective strategy for prioritizing wild relatives from germplasm banks for introgression into wheat. Continued reduction in the cost of genome sequencing and availability of reference genomes for the increasing number of wild relative species makes this strategy an attractive option for even large genebank collections including tens of thousands of lines (Mascher *et al*. 2019; Bohra *et al*. 2022). These approaches could quickly help to detect accessions enriched for alleles providing adaptation to target environments (Brunazzi *et al*. 2018; Anderson and Song 2020). As it was shown in our study, genome-wide introgression of prioritized diversity into adapted germplasm followed by fast high-throughput phenotyping using UAS-based imaging platforms could help to quickly identify promising germplasm for improving the adaptive potential of wheat. Expansion of these efforts from direct ancestors of bread wheat to include more distant *Aegilops* and *Triticum* species have potential to further broaden adaptive diversity accessible to wheat breeders for climate-proofing food production systems.

## Acknowledgements

This research was supported by the Agriculture and Food Research Initiative Competitive Grants 2022-68013-36439 (WheatCAP) and 2020-68013-30905 (USDA NIFA WWBI Hub) from the

USDA National Institute of Food and Agriculture.

## Author contribution

MN – data collection, analysis and interpretation, manuscript writing; DD – field data collection, UAS based phenotyping; EAd – population development, data analysis and field data collection; MC – population development and field data collection; HW – UAS imaging data analysis; AA – next-generation sequencing; AF – population development, data collection, experimental design; EAk – data analysis and interpretation, conceptualization, conceiving idea, manuscript writing.

## Supporting Information

**Fig. S1** The effect of geographic, climatic and bioclimatic variables on the SNP diversity in *Ae. tauschii* population.

**Fig. S2** Importance of canopy temperature and yield component traits for yield prediction based on random forest model in *Ae. tauschii*-Wheat introgression population.

**Table S1** Average historical climatic and bioclimatic data from the geographic origin of the diverse *Ae. tauschii*.

**Table S2** Number of climate associated SNPs for geographic climatic and bioclimatic variables discovered through redundancy analysis and genome-wide association analysis in *Ae. tauschii* population.

**Table S3** SNPs showing the highest correlation with geographic, climatic and bioclimatic variables in *Ae. tauschii* population.

**Table S4** Effect of SNPs that are significantly associated with geographic, climatic and bioclimatic variables in *Ae. tauschii* population based on genome-wide association analysis.

**Table S5** SNPs identified by redundancy analysis (RDA) and/or genome-wide association (GWAS) to be involved in *Ae. tauschii* ecogeographic adaptation.

**Table S6** High confidence climate associated SNPs identified by both redundancy analysis (RDA) and genome-wide assocition (GWAS) analysis in *Ae. tauschii* population.

**Table S7** SNPs significantly associated with canopy temperature in Ae. tauschii introgression population.

**File S1** Genome-wide association mapping of CAAs for adaptive traits in *Ae. tauschii*-wheat introgression population.

