## Supplementary Fig. 1 for "Ecogeographic signals of local adaptation in a wild relative help to identify variants associated with improved wheat performance under drought stress"

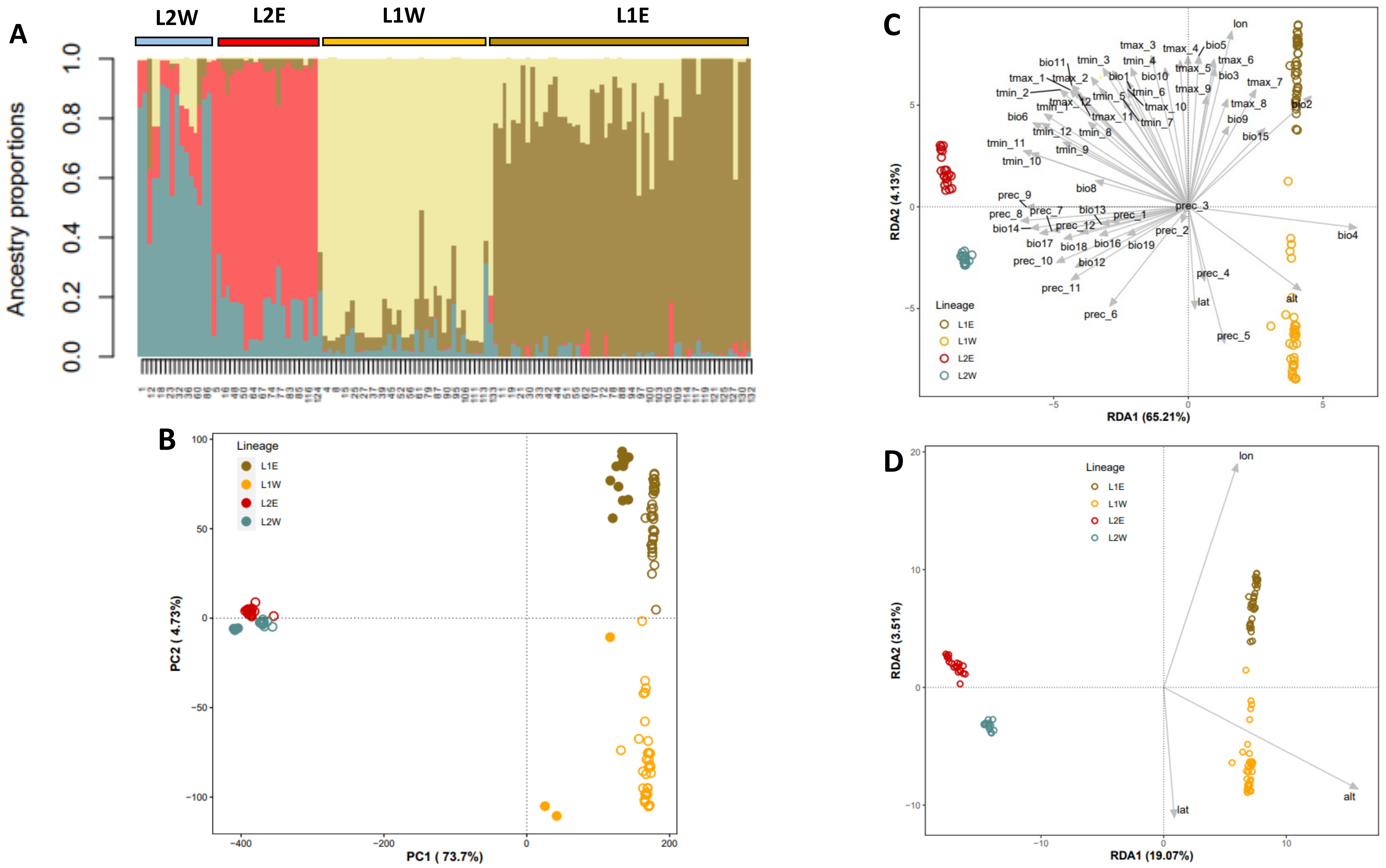


**Figure S1. The effect of geography, climatic and bioclimatic variables on the SNP diversity in *Ae. tauschii* population. A)** Stack barplot showing the ancestry proportion shared amongst the *Ae. tauschii* lines at K=4. **B)** Principal component plot for the 137 *Ae. tauschii* lines based on 109K SNPs. The open circles represent the 116 accessions whereas the filled circles represent the 21 lines used to generate the introgression population. **C)** Redundancy analysis biplot showing the effect of both geography and bioclimatic variables on *Ae. tauschii* diversity. Lineage 2 West (L2W) and Lineage 2 East (L2E) are *Ae. tauschii* ssp. *strangulata* whereas Lineage 1 East (L1E) and Lineage 1 West (L1W) are ssp. *tauschii*. **D)** Redundancy analysis biplot based on geographic variables
