## Supplementary File 1 for "Ecogeographic signals of local adaptation in a wild relative help to identify variants associated with improved wheat performance under drought stress"

**Heading date**

Heading date is one of the major adaptation traits that allowed wheat to colonize different agro-ecological zones globally (Cockram et al. 2007). It is controlled by several pathways that include vernalization (Vrn), photoperiod (*Ppd*) and gibberellin genes, which upon interaction with FLOWERING LOCUS C (FLC), induce floral initiation (Jung and Müller 2009). Fine tuning of the heading date in wheat has for long been considered a good strategy for improving wheat yield potential in specific environments (Reynolds et al. 2009; Hyles et al. 2020). Earliness per se (*Eps*) and Rht genes have been shown to play a role in fine tuning developmental phase transition in wheat thereby contributing to wheat adaptation (Lewis et al. 2008; Zikhali and Griffiths 2015; Basavaraddi et al. 2021a&b). The heading data for the introgression population lines were collected in 2020 at Ashland (AS20) and validated in 2022 at Rocky Ford (RF22), Kansas under rainfed conditions. At RF22, we had two replicate blocks whereas at Ashland we had one block from which heading data were collected. The number of days to heading (DTH) was calculated as the difference between heading date and planting date.

The correlation for DTH between the two blocks at RF22 was 0.8 whereas the correlation between the average DTH at RF22 and AS20 was 0.6. When the mixed linear model was fitted using the sommer package in R, the variance in DTH was not significantly affected by plot location within the block (Zratio = 0.0, P > 0.05) but the interaction between genotype and trial location (environment) was significant (Zratio = 3.337, P < 0.001) indicating the effect of genotype by environment interaction on heading date. Variation in heading date in the introgression population was calculated as the deviation of days to heading (DDTH) from the average days to heading for the control accessions. Based on the DDTH, some ILs headed five days earlier whereas others headed seven days later than the controls indicating that introgression of *Ae. tauschii* in hexaploid wheat introduced alleles that shifted heading date (Fig. 1A) in both directions. The early heading ILs (5% lower tail) were enriched with low frequency CAAs from *Ae. tauschii* whereas the late heading lines (5% upper tail) were enriched with low and high frequency CAAs (Fig. 3).

To understand the impact of heading date variation on yield, we compared the yield of ILs in the lower 5th and upper 95th percentiles of DDTH distribution using the data from non-irrigated trials in 2018-2020 because heading data were collected from non-irrigated trials. In 2018 and 2019 at Colby, the difference in yield was not significant between the groups (P = 0.98, P = 0.20, respectively), (Fig. 1C and 1D). However, in 2020, the late heading lines showed better yield compared to the early heading lines, P = 0.034 (Fig. 1E). Chen et al. (2018) showed that a dominant gibberellic acid-responsive dwarfing gene was associated with delayed heading date, increased spike fertility and harvest index. It is likely that a delay in heading allows the formation of more fertile spikelets. However, delayed heading is an undesirable trait but it can be reversed by the presence of *Ppd-D1a*, a photo-insensitive allele (Chen et al. 2018).

Based on the MLMM implemented in GAPIT3 (Segura et al. 2012; Wang and Zang 2021), we identified two genomic signals of association on chromosomes 2DS (chr2D_61772585) and 7DS (chr7D_35345332 and chr7D_55774657), (Fig. 1B). The alleles at chr2D_61772585 were C/G and the G allele from *Ae. tauschii* contributed to early heading in the introgression population. Similarly, the alleles at locus chr7D_35345332 and chr7D_55774657 were T/C and A/G,respectively. The alleles with reducing effect on heading date were C and G, and the G allele at chr7D_55774657 was observed in two hexaploid parents (KS061406LN_26 and Larry) and *Ae. tauschii* ssp. *strangulata* accessions. The C and A alleles from *Ae. tauschii* ssp. *strangulata* parents increased heading date. Two major genes *Ppd-D1* on 2DS and *VRN-D3* on 7DS are known to influence heading date in winter wheat (Wang et al. 2009). Variation of alleles at these loci in addition to other loci involved in flowering may explain why some introgression lines headed earlier than others relative to the controls (Fig. 1A).

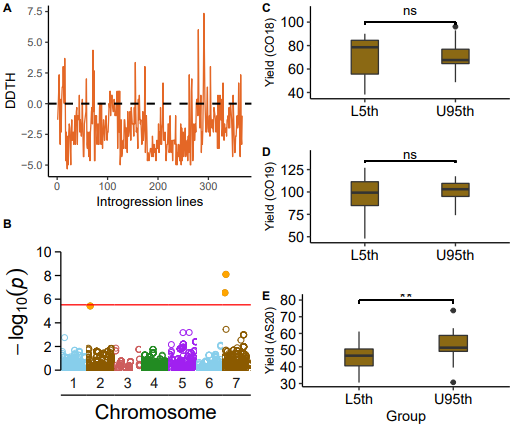

Figure 1. Relationship between the number of days to heading (DTH) and yield, and the genomic loci associated with days to heading of *Ae. tauschii* introgression lines. (A) Deviation in the number of days to heading (DDTH) for the ILs from the controls (parents and checks) mean days to heading. Yield (bushels per acre) for the ILs in the tails of DDTH distribution under non-irrigated conditions at Colby 2018 (B), Colby 2019 (C), and Ashland 2020 (D). Asterisks indicate significant difference (t-test, P =0.034). (E) Manhattan plot showing significant associations on 2DS and 7DS at a threshold FDR 0.05 indicated by the red line.

**Yield and yield components**

Yield improvement is one of the major goals for introgressing wild relative alleles in elite hexaploid wheat lines. It reflects the overall adaptive potential of the CAAs. Each CAA could have a small but additive effect on the final yield by positively contributing to one or a combination of the yield component traits. GWAS revealed that some of the CAAs on chromosomes 1DL, 2DS, 6DL and 7DS introgressed in hard red winter wheat were associated with yield and yield component traits such as SNS, grain width and length (Fig. 2). Alleles from *Ae. tauschii* had positive effects on SNS and GL but generally reduced GW. The phenotypic variance explained by the significant SNPs varied from 5-11% for the yield component traits. The significant SNPs on 2DS corresponded to the photoperiod response gene locus (*Ppd-D1*) known to influence heading date and spikelet number whereas the significant SNPs on 7DS are likely linked to *VRN-3D* which was shown to be involved in the regulation of developmental processes in wheat (Wang et al. 2009). Significant associations detected on 1DL and 6DL could possibly be linked to *Eps* genes which are known to act independently of *Pp-D1* and *VRN* genes to finetune the heading date and development phase transitions in wheat, further improving the adaptation of wheat to specific environments (Worland 1996; Lewis et al. 2008). Delayed heading also correlated with high number of spikelets per spike which could contribute to yield increase under favourable conditions. These results demonstrate that some of the CAAs introgressed in winter wheat have adaptive potential. Combining alleles that delay heading and increase spikelet number with alleles that confer drought and heat stress could mitigate the impact of climate change on wheat production.

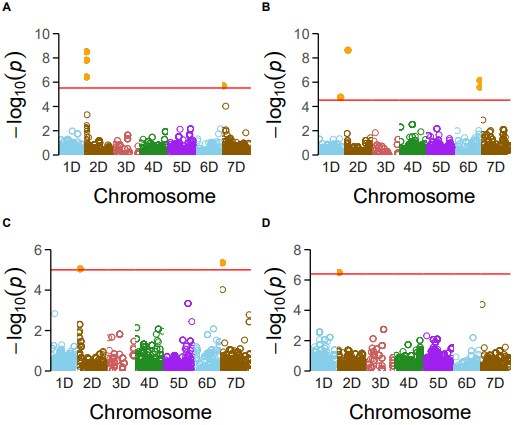

**Figure 2.** Manhattan plots showing CAAs on chromosomes 1D, 2D, 6D and 7D significantly associated with yield and yield component traits in the *Ae. tauschii* introgression population. Where (A) is grain length, (B) is grain width, (C) is spikelet number per spike and (D) is grain yield. Panel A-C are based on BLUPs as phenotypes and CMLM model whereas D is based on spatial adjusted yield at Colby in 2019 under non-irrigated conditions using MLMM model. SNPs above the red line are significantly associated with traits after FDR correction at alpha value 0.05.

| **SNP** | **Chr** | **Position** | **P.value** | **maf** | **nobs** | **Rsquare.of.**  **Model.**  **without.**  **SNP** | **Rsquare.of.**  **Model.with.**  **SNP** | **FDR_Adj_P-values** | **effect** | **Model** | **Trait** |
| --- | --- | --- | --- | --- | --- | --- | --- | --- | --- | --- | --- |
| chr2D_22255525 | 2 | 22255525 | 3.09E-09 | 0.26513 | 347 | 0.171733 | 0.263167 | 1.75E-05 | -0.084335578 | CMLM | Grain length (GL) |
| chr2D_22165390 | 2 | 22165390 | 1.55E-08 | 0.35879 | 347 | 0.171733 | 0.254621 | 4.39E-05 | 0.074587399 | CMLM | Grain length (GL) |
| chr2D_22276164 | 2 | 22276164 | 3.72E-07 | 0.304035 | 347 | 0.171733 | 0.238032 | 0.000704599 | -0.071871267 | CMLM | Grain length (GL) |
| chr7D_8728438 | 7 | 8728438 | 2.00E-06 | 0.279539 | 347 | 0.171733 | 0.229439 | 0.002838756 | 0.057702447 | CMLM | Grain length (GL) |
| chr2D_61772585 | 2 | 61772585 | 2.30E-09 | 0.253602 | 347 | 0.02615 | 0.135516 | 1.30E-05 | 0.029718751 | CMLM | Grain width (GW) |
| chr6D_462184586 | 6 | 462184586 | 7.11E-07 | 0.332853 | 347 | 0.02615 | 0.100202 | 0.002016761 | -0.023186001 | CMLM | Grain width (GW) |
| chr6D_462203291 | 6 | 462203291 | 2.62E-06 | 0.324207 | 347 | 0.02615 | 0.092403 | 0.004947538 | 0.022248565 | CMLM | Grain width (GW) |
| chr1D_432307193 | 1 | 432307193 | 1.73E-05 | 0.273775 | 347 | 0.02615 | 0.081253 | 0.016373083 | 0.022991084 | CMLM | Grain width (GW) |
| chr1D_432367341 | 1 | 432367341 | 1.73E-05 | 0.273775 | 347 | 0.02615 | 0.081253 | 0.016373083 | 0.022991084 | CMLM | Grain width (GW) |
| chr1D_432386924 | 1 | 432386924 | 1.73E-05 | 0.273775 | 347 | 0.02615 | 0.081253 | 0.016373083 | 0.022991084 | CMLM | Grain width (GW) |
| chr7D_13654672 | 7 | 13654672 | 4.45E-06 | 0.317771 | 332 | 0.168401 | 0.224891 | 0.025237531 | -0.242082536 | CMLM | Spikelet number per spike (SNS) |
| chr2D_31779118 | 2 | 31779118 | 8.93E-06 | 0.164157 | 332 | 0.168401 | 0.221202 | 0.025333185 | 0.259578291 | CMLM | Spikelet number per spike (SNS) |
| chr2D_22165390 | 2 | 22165390 | 3.21E-07 | 0.342988 | 328 | NA | NA | 0.001824437 | NA | MLMM | Yield |
| chr7D_55774657 | 7 | 55774657 | 8.00E-09 | 0.4121212 | 330 | NA | NA | 4.54E-05 | NA | MLMM | Days to heading (DTH) |
| chr7D_35345332 | 7 | 35345332 | 2.87E-07 | 0.075757 | 330 | NA | NA | 0.000815731 | NA | MLMM | Days to heading (DTH) |
| chr2D_61772585 | 2 | 61772585 | 3.76E-06 | 0.231818 | 330 | NA | NA | 0.007113905 | NA | MLMM | Days to heading (DTH) |

**Table 1.** SNPs significantly associated with days to heading, yield and component traits in *Ae. tasuchii* introgression population
