## Supplementary Table 2 for "Ecogeographic signals of local adaptation in a wild relative help to identify variants associated with improved wheat performance under drought stress"

**Table S2** Number of climate associated SNPs for geographic climatic and bioclimatic variables discovered through redundancy analysis and genome-wide association analysis in *Ae. tauschii* population.

| **Variable** | **Description** | **SNPs.RDA** | **SNPs.GWAS** |
| --- | --- | --- | --- |
| lon | Longitude | 2036 | 525 |
| lat | Latitude | 3 | 382 |
| alt | Altitude | 51 | 114 |
| bio1 | Annual Mean Temperature | 177 | 0 |
| bio2 | Mean Diurnal Range | 513 | 0 |
| bio3 | Isothermality | 18 | 0 |
| bio4 | Temperature Seasonality | 171 | 0 |
| bio5 | Max Temperature of Warmest Month | 5 | 134 |
| bio6 | Min Temperature of Coldest Month | 1184 | 1 |
| bio7 | Temperature Annual Range | 747 | 0 |
| bio8 | Mean Temperature of Wettest Quarter | 1 | 1 |
| bio9 | Mean Temperature of Driest Quarter | 889 | 555 |
| bio11 | Mean Temperature of Coldest Quarter | 173 | 0 |
| bio12 | Annual Precipitation | 4 | 1475 |
| bio13 | Precipitation of Wettest Month | 11 | 1976 |
| bio14 | Precipitation of Driest Month | 31 | 233 |
| bio15 | Precipitation Seasonality | 2 | 1050 |
| bio16 | Precipitation of Wettest Quarter | 0 | 1158 |
| bio17 | Precipitation of Driest Quarter | 7 | 227 |
| bio18 | Precipitation of Warmest Quarter | 7 | 351 |
| bio19 | Precipitation of Coldest Quarter | 1 | 0 |
| tmax_1 | Maximum Temperature in January | 1 | 36 |
| tmax_2 | Maximum Temperature in February | 19 | 66 |
| tmax_3 | Maximum Temperature in March | 54 | 40 |
| tmax_4 | Maximum Temperature in April | 20 | 0 |
| tmax_5 | Maximum Temperature in May | 2 | 105 |
| tmax_6 | Maximum Temperature in June | 76 | 113 |
| tmax_7 | Maximum Temperature in July | 18 | 82 |
| tmax_8 | Maximum Temperature in August | 14 | 82 |
| tmax_9 | Maximum Temperature in September | 0 | 82 |
| tmax_10 | Maximum Temperature in October | 0 | 65 |
| tmax_11 | Maximum Temperature in November | 2 | 0 |
| tmax_12 | Maximum Temperature in December | 41 | 73 |
| tmin_1 | Minimum Temperature in January | 2 | 0 |
| tmin_2 | Minimum Temperature in February | 190 | 36 |
| tmin_3 | Minimum Temperature in March | 702 | 38 |
| tmin_4 | Minimum Temperature in April | 58 | 220 |
| tmin_6 | Minimum Temperature in June | 9 | 0 |
| tmin_7 | Minimum Temperature in July | 0 | 20 |
| tmin_8 | Minimum Temperature in August | 2 | 360 |
| tmin_9 | Minimum Temperature in September | 7 | 30 |
| tmin_10 | Minimum Temperature in October | 83 | 0 |
| tmin_11 | Minimum Temperature in November | 337 | 0 |
| tmin_12 | Minimum Temperature in December | 32 | 49 |
| prec_1 | Precipitation in January | 1 | 943 |
| prec_2 | Precipitation in February | 3 | 155 |
| prec_3 | Precipitation in March | 4 | 111 |
| prec_4 | Precipitation in April | 0 | 179 |
| prec_5 | Precipitation in May | 6 | 0 |
| prec_6 | Precipitation in June | 3 | 767 |
| prec_7 | Precipitation in July | 38 | 617 |
| prec_8 | Precipitation in August | 275 | 5073 |
| prec_9 | Precipitation in September | 2085 | 6362 |
| prec_10 | Precipitation in October | 0 | 5866 |
| prec_11 | Precipitation in November | 13 | 3516 |
| prec_12 | Precipitation in December | 21 | 109 |
