## Supplementary figures and images for "Ecogeographic signals of local adaptation in a wild relative help to identify variants associated with improved wheat performance under drought stress"

### Supplementary Figure 2

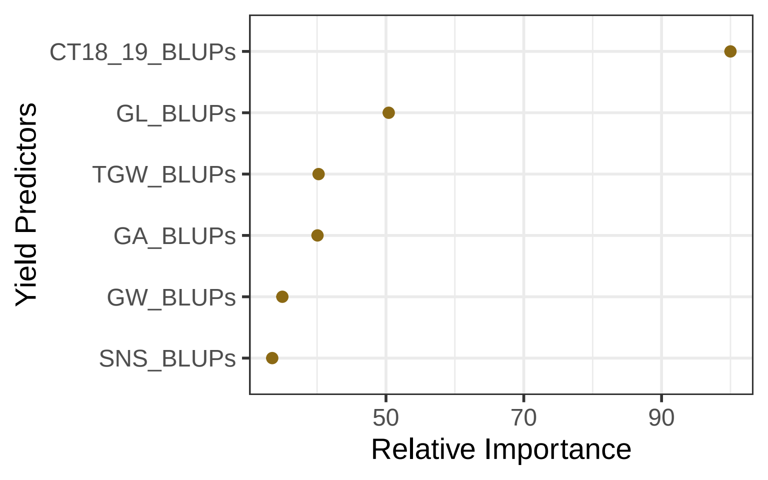
